# Aminoglycoside antibiotics inhibit mycobacteriophage infection

**DOI:** 10.1101/2020.07.03.185546

**Authors:** Zheng Jiang, Junwei Wei, Nan Peng, Yingjun Li

## Abstract

Antibiotic resistance is becoming the biggest current threat to global health. At the same time, phage therapy is witnessing a return of interest. The therapeutic use of bacteriophages, that infect and kill bacteria, is well suited to be a good strategy to combat antibiotic resistance. Furthermore, bacteriophages are increasingly used in combination with standard antibiotics against the drug-resistant pathogens. Interestingly, we found that the engineered mycobacteriophage phAE159 and natural phage D29 can not infect the *Mycobacterium tuberculosis* in the presence of kanamycin, hygromycin or streptomycin, but there is no effect on the phage infection in the presence of spectinomycin. Based on a series of studies and structural analysis of the above four aminoglycoside antibiotics, we can speculate as to the mechanism by which amino sugar group of aminoglycoside was able to selectively inhibit mycobacteriophage DNA replication. This is a rare discovery that broad-spectrum antibiotics inhibit phage infection. We envisioned that this study will provide guidance for people to combine phage and antibiotics to treat *M. tuberculosis*.

## INTRODUCTION

Bacterial infection refers to the invasion of a host’s tissue by pathogenic bacteria. Generally, antibiotics are the preferred antibacterial agents 1. However, bacteria can evolve resistance to antibiotics resulting from antibiotics abusage and natural evolution. Bacteriophage provides a alternative useful antibacterial approach, and has gradually been used in combination with standard antibiotics against the drug-resistance of pathogenic bacteria 2. An important question arises: Do antibiotics impact the pharmacodynamics of phage therapy?

In this study, we firstly found that the replication of mycobacteriophage was significantly inhibited by aminoglycoside antibiotics. We speculated as to the mechanism by which amino sugar group of aminoglycoside was able to selectively inhibit mycobacteriophage DNA replication. The discovery in this study may shake the previous understanding of the synergy of the phage and antibiotic therapies, providing guidance for people to combine phage and antibiotics to treat *M. tuberculosis*.

## MATERIALS AND METHODS

### Transduction of mycobacteriophage plasmid

*M. smegmatis* mc^2^155::pMV261 was grown in 7H9 to an OD_600_ of 1.0(~6×10^8^ c.f.u. mL^−1^). Hundred milliliters of the culture was centrifuged and washed 3 times by 10% glycerol and resuspended in 5 ml 10 % glycerol. The phAE159 plasmid were electransformed into *M*. *smegmatis* mc^2^155. Cells were mixed with 3 mL of 7H9 top agar (containing 0.75% agar) and with/without 50 μg/mL kanamycin, then plated on 7H10 agar plates with/without 50 μg/mL kanamycin, and then incubated at 30°C for 3 days. The number of visible plaques was counted. Both 7H9 agar and 7H10 agar were added when the temperature of the medium was below 55°C.

### Phage propagation assay

The phage D29 and phAE159 was collected and serial dilutions by MP buffer, then they were spotted onto the lawns of *Mycobcaterium*. Plates were incubated at 37 °C (30 °C for phAE159) overnight and phages were enumerated by counting plaques. These assays were repeated at least three times with similar results and a representative experiment is shown.

### Construction of pSTR1

The *aadA* gene was amplified from pCDFDuet-1 plasmid using the primers STR-F and STR-R in Table S3 and cloned into the pMV261 vector. The plasmid was transformed into *M. smegmatis* mc^2^155 by standard procedures 3. The recombinant strains were selected on 7H10 agar plates complemented with 50 μg/mL streptomycin and 50 μg/mL kanamycin. The positive clone was named mc^2^155::pSTR1. Then the mc^2^155::pSTR1 were incubated in 7H10 agar plates complemented with 50 μg/mL streptomycin or 100 μg/mL spectinomycin.

### Growth curves analysis of *E. coli* being infected by phage T7

*E. coli* strain DH5α was grown overnight with shaking in lysogeny broth (LB) medium at 37 °C. A 2% subculture was prepared in LB medium supplemented with kanamycin (50 μg/mL) and phage T7 was added at a multiplicity of infection of approximately 0.1. Subsequently, the number of DH5α present in cultures was determined by optical density at 600 nm every 2 h for an additional 15 h. Three biological replicates were tested for each group of kanamycin as well as the control, which contained no kanamycin or phage T7.

### Pre-incubation of phage phAE159 with kanamycin

Aliquots of phage phAE159 were incubated at 37 °C for 2 h with or without 50 μg/mL kanamycin. The phages were diluted with MP buffer and added into *M. smegmatis* mc^2^155::pMV261 cells, then the mixture was suspended in molten 3 mL of 7H9 top agar (with/without 50 μg/mL kanamycin) and overlaid onto 7H10 plates pre-added with kanamycin or not. After incubating the plates at 30 °Cfor 2 days, observe the growth of *M. smegmatis* mc^2^155.

### Transmission electron micrograph analysis

*M. smegmatis* mc^2^155 was grown in 7H9 medium at 37 °C with shaking, and the cells were harvested until the OD_600_ (optical density at 600 nM) = 0.8. High titer phage D29 were then added into *M. smegmatis* mc^2^155 at a multiplicity of infection (MOI) of 100. The culture was incubated for 15 minutes with/without 50 μg/mL streptomycin before microscopic observation. Transmission electron microscopy (TEM) grids (Electron Microscopy Sciences CF400-CU) were prepared by drop-coating grids with each group, washing with water and staining with 2% uranyl acetate. Phages were imaged using the HITACHI H-7650 transmission electron microscope in the Microscopy Imaging Laboratory at the Huazhong Agricultural University.

### Quantitative PCRs analysis

*M. smegmatis* mc^2^155::pMV261 was grown in 7H9 with 10 folds phage D29 and 50 μg/mL streptomycin. Sampling culture fluid and centrifuged supernatant every 1 hour. Quantitative PCR (qPCR) was performed as standard procedures to quantify D29-DNA 4. Each reaction mixture (20 μL/well) contained 10 μL of Hieff qPCR SYBR Green Master Mix No Rox, 1 μL of 10 μM each of the gp69-qpcr-F and gp69-qpcr-R primer pair in Table S3, 7 μL of DNase- and RNase-free sterile water, and 1 μL of the sample or template DNA. Quantitative PCR and monitoring were performed in an ABI Quant Studio 5 System. PCR amplification was performed with an initial pre-incubation at 95◦C for 5 min, followed by 40 cycles of amplification at 95 °C for 10 s, 60 °C for 20 s and 72 °C for 20 s. A 810 bp PCR product was generated using the standard PCR primers gp69-F/R in Table S3 and D29 DNA to construct the standard curve. After purification and determination of the DNA concentration, the linear double-stranded DNA standard was 10-fold serially diluted to obtain a standard series from 1×10^7^ to 1×10^1^ copies/μL. The copy numbers of the samples were determined by reading off the standard series with the Ct values of the samples.

## RESULTS AND DISCUSSION

Our primary goal was to eliminate *Mycobacterium tuberculosis* by an engineered mycobacteriophage (unpublished data). In order to avoid the contamination of microbes in infection test, we used a *Mycobacterium tuberculosis* strain that carrying a plasmid with kanamycin-resistance gene and added kanamycin in the culture. Interestingly, we found that the engineered mycobacteriophage can not infect the host strain. Then, we examined the ability of kanamycin to inhibit the infection of the TM4-derived phasmid phAE159 5 and D29. In the absence of kanamycin, the two phages were able to form plaques on lawns of *M. smegmatis* mc^2^155. In the presence of the above antibiotics, the replication of the two mycobacteriophages was inhibited 10^3^-fold or more (Fig. 1a). We determined an inhibitory concentration of 50 μg/mL for kanamycin inhibiting phage D29, which does not affect the growth of *M. smegmatis* mc^2^155 (Fig. S1). Furthermore, we verified this effect in the infection test of *M. tuberculosis* H37Ra and *M. bovis* BCG by phage D29 (Fig. S2). This demonstrates that kanamycin can suppress the replication of the mycobacteriophages phAE159 and D29.

**Fig. 1.**
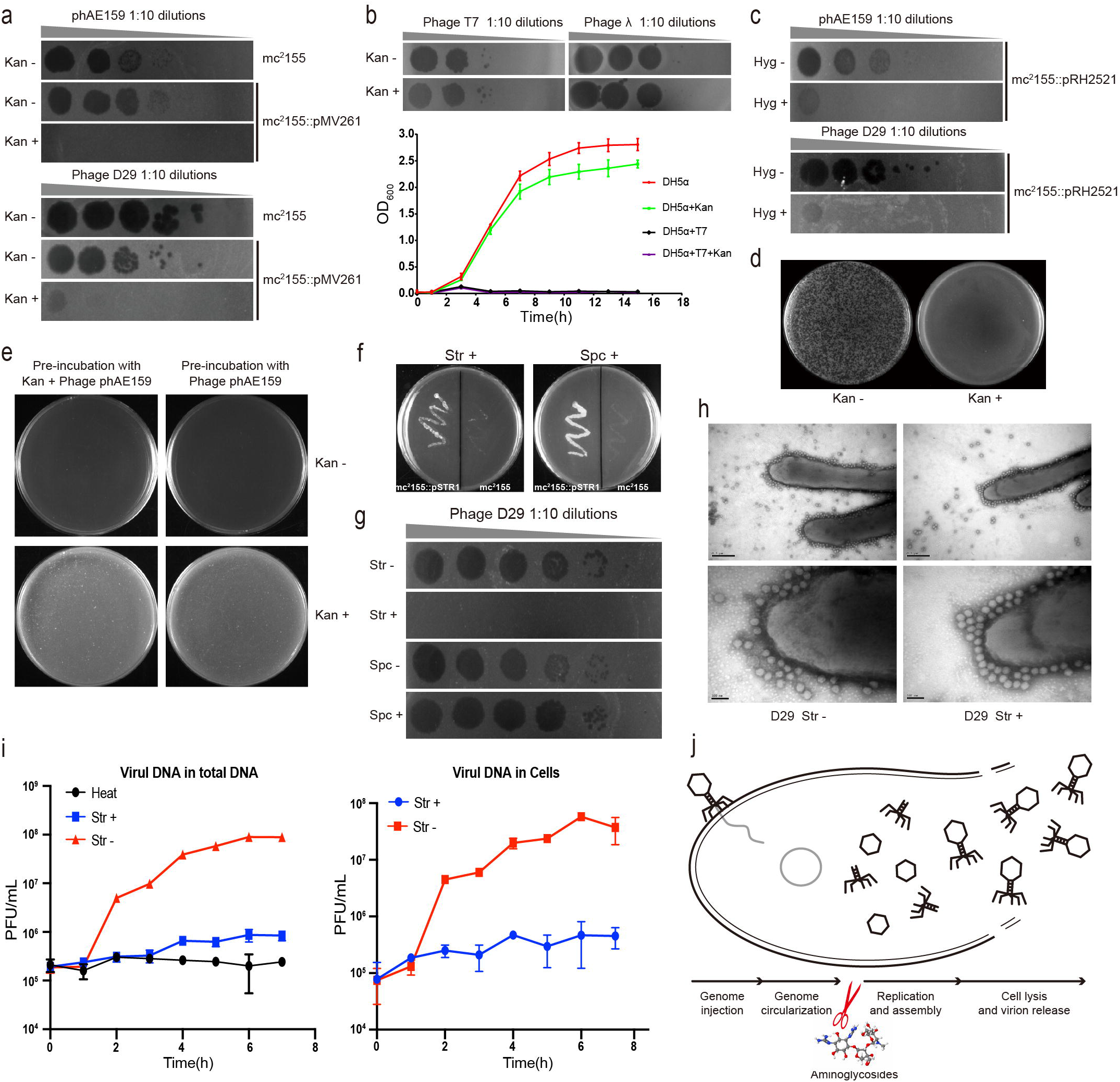
Aminoglycoside antibiotics inhibit the DNA replication of mycobacteriophages. **a.** The replication of the two mycobacteriophages phAE159 and D29 were inhibited by kanamycin on the lawns of *M. smegmatis* mc^2^155. **b.** Kanamycin could not affect the infection of *Escherichia coli* phages T7 and λ. **c.** Hygromycin was able to inhibit the infection of mycobacteriophages phAE159 and D29. **d.** Kanamycin inhibits the formation of plaques by electrotransformation of phAE159 vector. **e.** Pre-incubation with kanamycin does not affect the infection of mycobacteriophage phAE159. **f.***M. smegmatis* mc^2^155 harboring the recombinant plasmid tolerated streptomycin and spectinomycin well. **g.** The propagation of mycobacteriophage D29 was significantly inhibited in the presence of streptomycin but had no effect in the presence of spectinomycin. **h.** TEM analysis of the adsorption capacity of mycobacteriophage in the presence of streptomycin. **i.** Quantitative PCR analysis of phage DNA proliferation during the infection process. **j.** Aminoglycosides were speculated to be able to block the DNA replication during the life cycle of phage (adapted from Fig.3a in 6).

To determine whether other bacteriophages also can be suppressed by kanamycin, we examined the ability of kanamycin to protect *Escherichia coli* from lysis by the well-characterized dsDNA phages T7 and λ. In the plaque and growth curve assays, we found that the presence of kanamycin did not affect the infection of *E. coli* phages (Fig. 1b). It suggested that this inhibitory effect might be specific to mycobacteriophages. In addition, we examined this effect for another commonly used antibiotic in *Mycobacterium*, hygromycin, which is also an aminoglycoside antibiotic. It was observed that the hygromycin was able to inhibit mycobacteriophages phAE159 and D29 effectively (Fig. 1c). Following, we examined the ability of kanamycin to suppress the lysis of *M. smegmatis* by the electroporation of the mycobacteriophage vector phAE159. The plaques were observed in the absence but not in the presence of kanamycin (Fig. 1d).

In order to study whether the antibiotic directly acts on the phages, e.g., destroys the phages. *M. smegmatis* mc^2^155 were incubated with mycobacteriophages phAE159 and with/out kanamycin, then plated on 7H10 with/out kanamycin. The lawn of each treatment that are present after 2 days growth of *M. smegmatis* showed that regardless of whether kanamycin was added during pre-incubation, as long as the kanamycin is in the solid medium in plates, the bacteria will not be lysed by the phage and grow well (Fig. 1e).

The two antibiotics we used earlier are aminoglycosides. Therefore, we considered whether this inhibitory effect is applicable to other aminoglycoside antibiotics. First, we constructed a plasmid pSTR1 carrying *aadA* (encoding aminoglycoside adenylyltransferase) gene, which confers resistance to streptomycin and spectinomycin. The antibiotic sensitivity testing experiments showed that *M. smegmatis* mc^2^155 harboring the recombinant plasmid tolerated streptomycin and spectinomycin well, while the WT could not grow in the presence of either of these two antibiotics (Fig. 1f). Next, we used propagation assays to explore the effect of these two antibiotics on the infection of mycobacteriophages. We found that the propagation of phage on the plate was significantly inhibited in the presence of streptomycin but had no effect in the presence of spectinomycin (Fig. 1g).

To explore the mechanism more intuitively, transmission electron microscope (TEM) analysis was performed to determine whether the adsorption capacity of mycobacteriophage is normal in the presence of antibiotics. Before microscopic observation, we mixed phage D29 and *M. smegmatis* mc^2^155 for 15 mins. We can see that the addition of streptomycin has no effect on the phage morphology, indicating that streptomycin cannot directly act on the phage. And phages can also attach to the surface of the bacteria normally, there is no difference compared with the control group (Fig. 1h). This indicates that streptomycin does not inhibit the adsorption of mycobacteriophage to the cell surface of *Mycobacterium*. After the bacteriophage attached to the surface of the bacteria, the injection and cyclization of its genome DNA is a very rapid process. Therefore, we speculate that the inhibitory effect of antibiotics on phage infection may act after viral genome injection. We next used absolute quantification PCR to detect the content of phage DNA during the infection process. Bacteria were cultured in the presence of streptomycin, mycobacteriophages were then added at a high multiplicity of infection and detect the content of phage DNA in culture and supernatant every hour. In the absence of streptomycin, phage DNA increased exponentially after 1 hour of co-culture, While the phage DNA did not grow in the presence of streptomycin. This phenomenon was also observed for phage DNA in the host cells, indicating that streptomycin inhibits mycobacteriophage infecting host by block phage DNA replication (Fig. 1i).

The life cycle of a phage starts with the adsorption and injection of the genome into the host cell. After DNA injection, the phage genome circularizes, then the DNA replicates and assembles, finally the host cell is lysed, and the progeny phages are released. In briefly summary, in this study, visually observation of the morphology by TEM indicated that the aminoglycoside antibiotics does not prevent the phage adsorbing the host cell. The electroporation and pre-incubation of mycobacteriophage phAE159 and *M. smegmatis* mc^2^155 with/out kanamycin showed that the inhibition of kanamycin on phage occurs after the phage injects DNA into the host cell. Moreover, we observed that the mycobacteriophage DNA could not proliferate in host cells in the presence of streptomycin. Taken together, we determined that aminoglycosides were able to block phage DNA replication and allow the bacteria to survive (Fig. 1j).

Although the antibiotics we used here are all classified as aminoglycosides because they all contain amincyclic alcohols, spectinomycin differs from the other three antibiotics in that it does not have an amino sugar group (Fig. S3). Based on the above observations and results, at this time, we can only speculate as to the mechanism by which amino sugar group in aminoglycoside was able to selectively inhibit mycobacteriophage DNA replication. Therefore, further investigations to explore their mechanisms of action will expand our knowledge of bacterial anti-phage defense systems 6 and the arms race between bacteria and phage foes 7. In addition, it is particularly noteworthy that streptomycin is a first-line drug for tuberculosis, thus the discovery in this study may shake the previous understanding of the synergy of the two therapies, phages and antibiotics 8. We envisioned that this study will provide guidance for people to combine phage and antibiotics to treat *M. tuberculosis*.

## ACKNOWLEDGMENTS

We are grateful to Prof Chen Tan, Prof Jinsong Li, Prof Hui Jin and Yibao Chen for providing phAE159, D29, λ and T7 phages, respectively. We also thank Dr Min Yang and Dr Hua Zhang for providing *Mycobacterium* strains and the shuttle vector pMV261. This work was supported by the National Natural Science foundation of China (31801035), National Postdoctoral Program for Innovative Talents (BX201800113), Natural Science Foundation of Hubei Province (2018CFB223), China Postdoctoral Science Foundation (2018M640708).

## Author contributions

ZJ, JW, NP and YL designed experiments. ZJ, JW and YL performed the experiments. ZJ, JW, NP and YL analyzed the data presented in the manuscript. All authors discussed and approved the manuscript.

## Conflicts of interest

No potential conflicts of interest were disclosed.

## Notes

### Competing Interest Statement

The authors have declared no competing interest.

