## Supplemental information for "Aminoglycoside antibiotics inhibit mycobacteriophage infection"

### Supplementary tables

**Table S1. Strains and phages used in this work**

| Strains and Phages | Reference |
|---|---|
| *M. tuberculosis* H37Ra | Gift from Dr Min Yang |
| *M. bovis* BCG | Gift from Dr Min Yang |
| *M. smegmatis* $mc^2155$ | Gift from Dr Min Yang |
| *E. coli* strain DH5α | Laboratory stock |
| Mycobacteriophage D29 | Gift from Prof Jinsong Li |
| *E. coli* phage T7 | Gift from Yibao Chen |
| *E. coli* phage λ | Gift from Prof Hui Jin |

**Table S2. Plasmids used in this work**

| Plasmids | Genotype and features | Reference |
|---|---|---|
| pMV261 | A *Mycobacterium-E. coli* shuttle vector carrying kanamycin resistance gene | Gift from Dr Hua Zhang |
| pRH2521 | A *Mycobacterium-E. coli* shuttle vector carrying hygromycin resistance gene | Gift from Dr Min Yang |
| pSTR1 | Derived from pMV261，carrying aminoglycoside adenylyltransferase, confers resistance to streptomycin and spectinomycin | This work |
| pET28a | A *E. coli* expression vector carrying kanamycin resistance gene | Laboratory stock |
| phAE159 | A mycobacteriophage vector derived from TM4 | (1), Gift from Prof Chen Tan |

**Table S3. Oligonucleotides used in this work**

| Oligonucleotide | Sequence (5’-3’) |
|---|---|
| STR-F | GCAATGGCCAAGACAATTGCGGATATGAGGGAAGCGGTGATCG |
| STR-R | GCCTGCTGATGATGTCTTAATTAAGGATCTTATTTGCCGACTACCTTGGTGA |
| gp69-qpcr-F | AGACCGGCGACTACTTCATGG |
| gp69-qpcr-R | GCAACGGGTCGAACATCGAG |
| gp69--F | GTGACGCAGATCAAGCTTCC |
| gp69--R | TCACTTAAAAACGGGGCAACTG |

### Supplementary figures

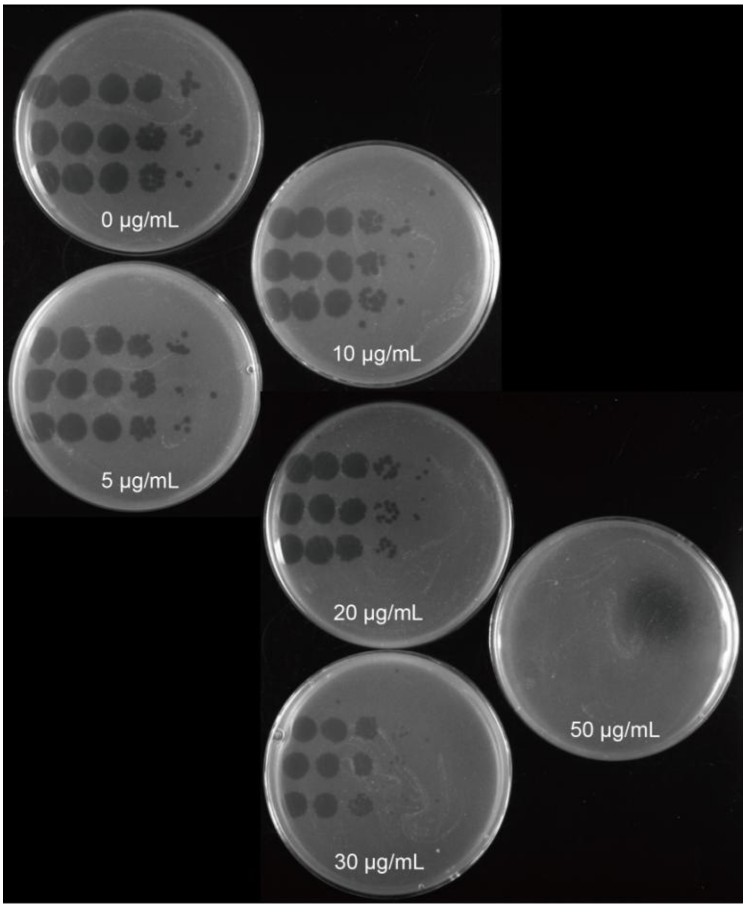


**Fig. S1. Effects of different concentrations of kanamycin on the infection of *M. smegmatis* mc$^2$155 by phage D29**

Phages were gradually 10-fold diluted from left to right, three replicates on each plate and three biological replicates were performed.

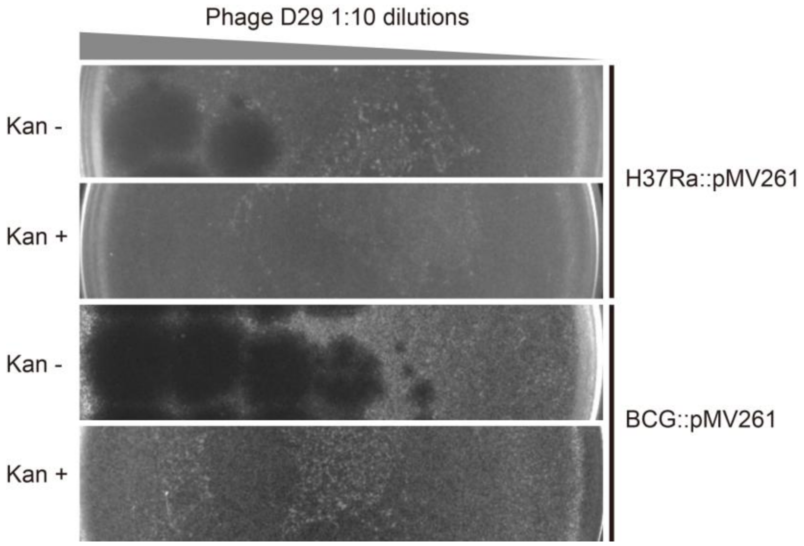


**Fig. S2. Kanamycin inhibit the infection of *M. tuberculosis* H37Ra and *M. bovis* BCG by phage D29**

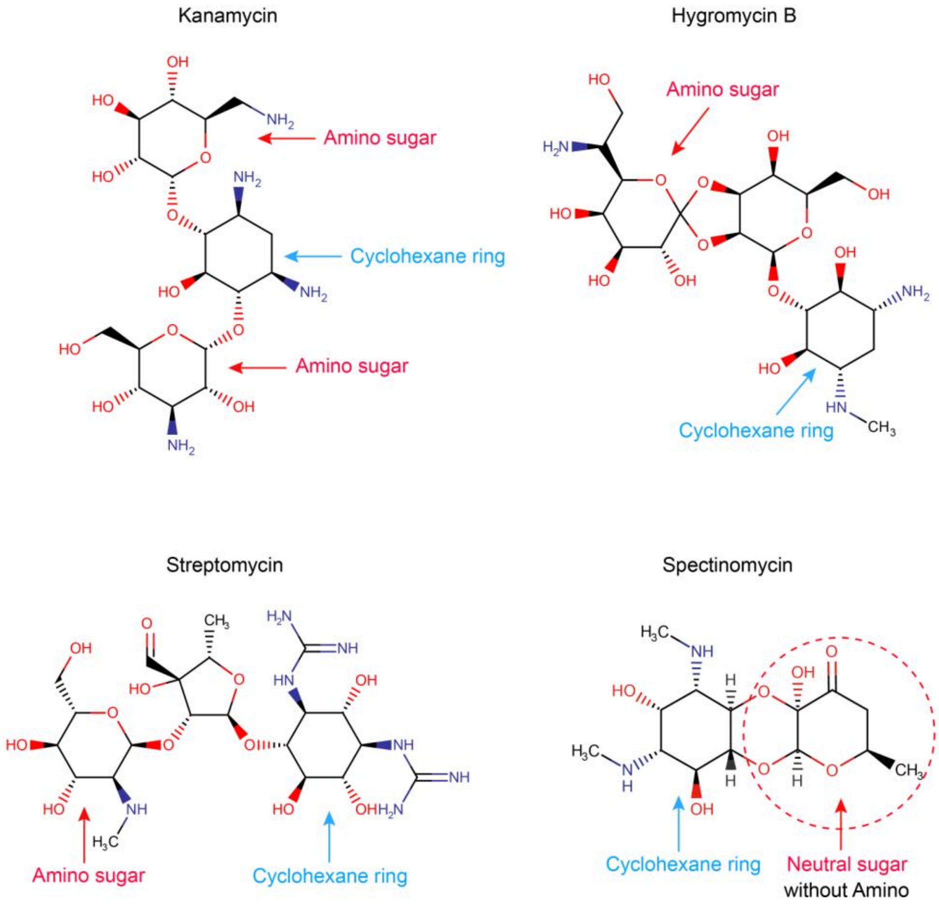


**Fig. S3. The structures of the four aminoglycoside antibiotics used in this study**

Amino units are in Blue, and Cyclohexane ring and amino sugar are marked with cyan-blue and red arrows, respectively. The neutral sugar without amino in spectinomycin is marked with red dotted circle.
